## Supplementary material for "Multi-locus genotyping reveals established endemicity of a geographically distinct *Plasmodium vivax* population in Mauritania, West Africa": S1 Table

Supplementary Table S1. List of 38 *P. vivax* SNP marker loci analysed in this study.

| Marker (Chr:SNP position) | Reference Allele | Alternative Allele |
| --- | --- | --- |
| PvP01_01_v1:612295 | T | C |
| PvP01_01_v1:778121 | T | G |
| PvP01_02_v1:527792 | T | C |
| PvP01_03_v1:239662 | T | C |
| PvP01_03_v1:574236 | G | A |
| PvP01_03_v1:595569 | A | G |
| PvP01_04_v1:367674 | C | T |
| PvP01_05_v1:163535 | A | G |
| PvP01_05_v1:1409127 | C | T |
| PvP01_06_v1:76255 | A | G |
| PvP01_06_v1:602491 | C | T |
| PvP01_08_v1:173904 | C | T |
| PvP01_08_v1:577308 | G | A |
| PvP01_08_v1:1419576 | C | T |
| PvP01_08_v1:1565446 | C | T |
| PvP01_09_v1:846288 | T | C |
| PvP01_10_v1:151400 | T | G |
| PvP01_10_v1:865053 | T | G |
| PvP01_10_v1:1304923 | T | C |
| PvP01_10_v1:1371780 | C | T |
| PvP01_11_v1:108200 | T | C |
| PvP01_11_v1:256884 | T | C |
| PvP01_11_v1:643929 | C | T |
| PvP01_11_v1:747140 | A | G |
| PvP01_11_v1:1809904 | G | A |
| PvP01_11_v1:1969060 | A | G |
| PvP01_12_v1:1139454 | T | C |
| PvP01_12_v1:1220748 | C | T |
| PvP01_12_v1:2685195 | G | A |
| PvP01_13_v1:507029 | C | T |
| PvP01_13_v1:851321 | A | G |
| PvP01_13_v1:1088863 | T | C |
| PvP01_13_v1:1160640 | C | A |
| PvP01_14_v1:439002 | G | A |
| PvP01_14_v1:855554 | T | C |
| PvP01_14_v1:1036358 | C | T |
| PvP01_14_v1:2258311 | A | G |
| PvP01_14_v1:2310569 | C | A |

SNP positions are based on the PvP01 *P. vivax* reference genome sequence (Auburn et al. 2016 Wellcome Open Res 1:4 <https://doi.org/10.12688/wellcomeopenres.9876.1>)
