## Supplementary material for "Multi-locus genotyping reveals established endemicity of a geographically distinct *Plasmodium vivax* population in Mauritania, West Africa": S2 Table

**Supplementary Table S2.** Multi-locus linkage disequilibrium in *P. vivax* in Mauritania and in previous data from other countries.

|  | **All infections** |  | **Unique haplotypes** |  |
| --- | --- | --- | --- | --- |
| **Site** | ***N*** | ***I*_A_^S^** | ***N*** | ***I*_A_^S^** |
| Nouakchott | 87 | 0.0024 ^NS^ | 85 | 0.0021 ^NS^ |
| Zouerat | 6 | -0.0118 ^NS^ | 6 | -0.0118 ^NS^ |
| Mauritania | 94 | 0.0022 ^NS^ | 91 | 0.0022 ^NS^ |
| Ethiopia | 22 | 0.0045 ^NS^ | 21 | 0.0022 ^NS^ |
| Thailand | 100 | 0.0007 ^NS^ | 96 | 0.0006 ^NS^ |
| Indonesia | 104 | 0.0026 ^NS^ | 102 | 0.0022 ^NS^ |
| Mexico | 19 | 0.0857 *** | 16 | 0.0882 *** |
| Colombia | 30 | 0.0173 *** | 26 | 0.0114 ** |

The standardised Index of Association (***I*_A_^S^** ) tests for departures from multi-locus equilibrium, with significant values being above zero as indicated: ^NS^ Not significant (*P* > 0.05), * *P* < 0.05, ** *P* < 0.01, *** *P* < 0.001.

Data are as described in the Methods and Results for the array of 38 SNPs, with only monoclonal infections having complete genotypes at 37 SNPs (excluding PvP01_03_v1:595569) being included in this analysis.
