## Supplementary material for "Multi-locus genotyping reveals established endemicity of a geographically distinct *Plasmodium vivax* population in Mauritania, West Africa": S3 Table

**Supplementary Table S3.** Genetic differentiation between the Mauritanian *P. vivax* population sampled in this study and previously described populations.

|  | **Mauritania** | **Ethiopia** | **Thailand** | **Indonesia** | **Mexico** | **Colombia** |
| --- | --- | --- | --- | --- | --- | --- |
| **Mauritania** | - | 0.14 | 0.18 | 0.20 | 0.17 | 0.16 |
| **Ethiopia** | 0.22 | - | 0.14 | 0.22 | 0.25 | 0.14 |
| **Thailand** | 0.28 | 0.27 | - | 0.13 | 0.2 | 0.13 |
| **Indonesia** | 0.31 | 0.32 | 0.21 | - | 0.3 | 0.25 |
| **Mexico** | 0.29 | 0.33 | 0.39 | 0.44 | - | 0.11 |
| **Colombia** | 0.23 | 0.2 | 0.21 | 0.33 | 0.17 | - |

Hudson’s *F*_ST_ index is shown in the lower matrix, Weir & Cockerham’s *F*_ST_ index in the upper matrix, each based on the mean of the array of 38 SNPs with data as described in the Methods and Results.
