## Supplementary figures and images for "Multi-locus genotyping reveals established endemicity of a geographically distinct *Plasmodium vivax* population in Mauritania, West Africa"

### S1 Fig

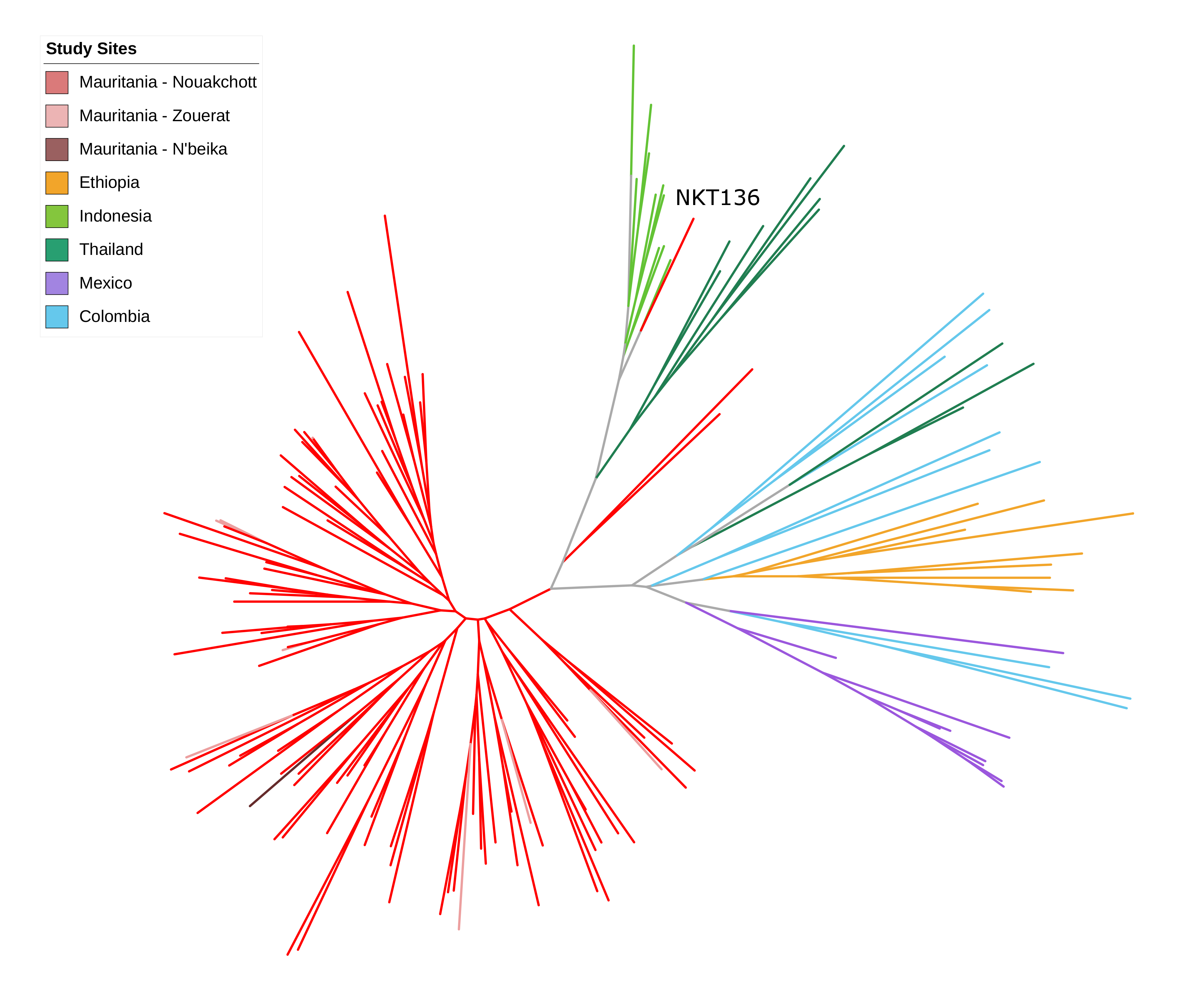

### S2 Fig

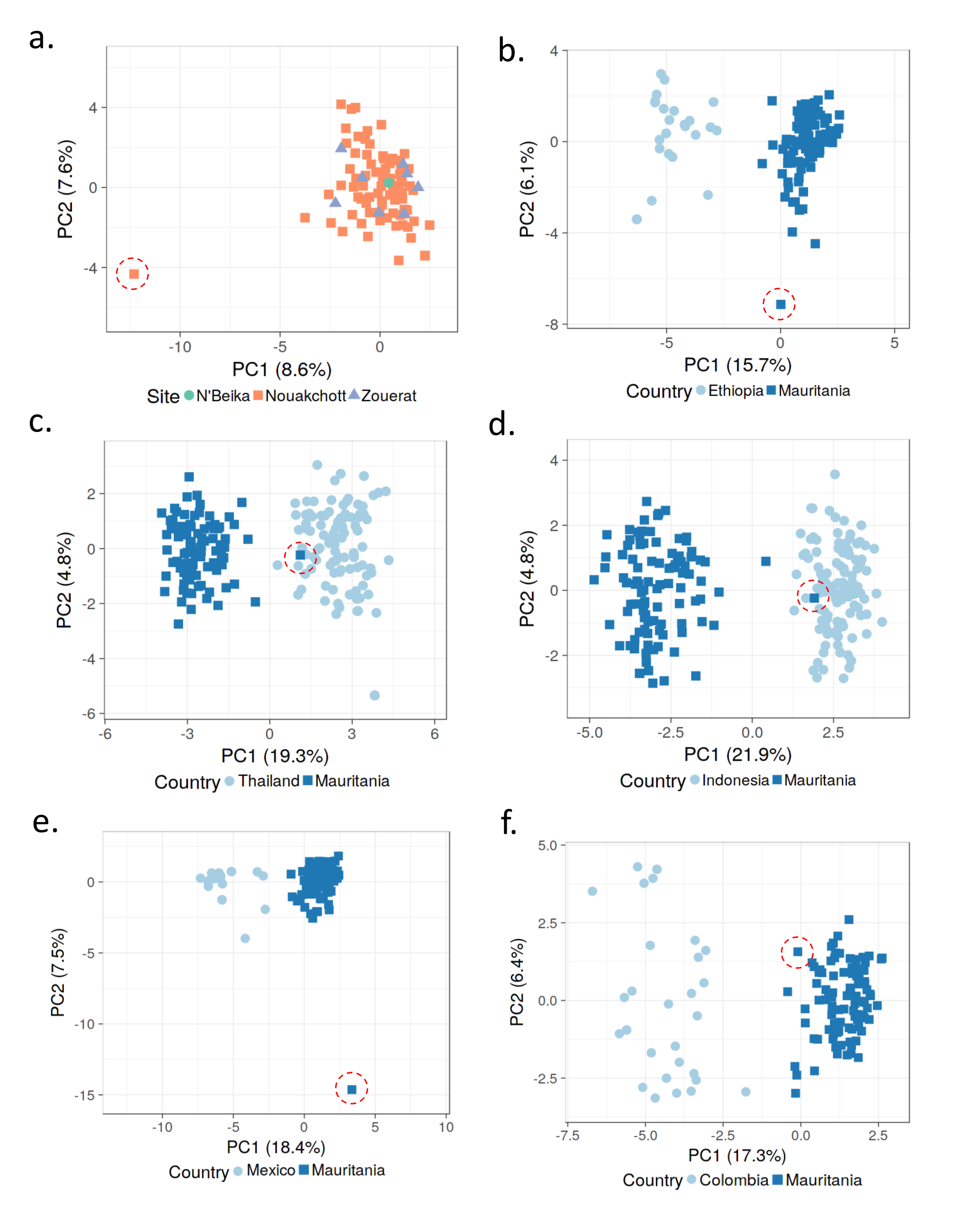
